## supplemental notes, figures, and tables for "StratoMod: Predicting sequencing and variant calling errors with interpretable machine learning"

#### Supplemental Note 1

The following was used to generate the gene counts for **Supplemental File 1.**

```
intersectBed -c -a GRCh38_mrg_full_gene.bed -b patho_hifi_ilm.tsv > \
    GRCh38_mrg_full_gene_FNcounts.bed

grep Illumina patho_hifi_ilm.tsv | \
    intersectBed -c -a GRCh38_mrg_full_gene.bed -b stdin > \
    GRCh38_mrg_full_gene_FNcountsIllonly.bed

grep both patho_hifi_ilm.tsv | \
    intersectBed -c -a GRCh38_mrg_full_gene.bed -b stdin > \
    GRCh38_mrg_full_gene_FNcountsboth.bed

grep SNV patho_hifi_ilm.tsv > patho_hifi_ilm_SNV.tsv

intersectBed -c -a GRCh38_mrg_full_gene.bed -b patho_hifi_ilm_SNV.tsv > \
    GRCh38_mrg_full_gene_SNV_FNcounts.bed

grep Illumina patho_hifi_ilm_SNV.tsv | \
    intersectBed -c -a GRCh38_mrg_full_gene.bed -b stdin \
    > GRCh38_mrg_full_gene_SNV_FNcountsIllonly.bed

grep both patho_hifi_ilm_SNV.tsv | \
    intersectBed -c -a GRCh38_mrg_full_gene.bed -b stdin > \
    GRCh38_mrg_full_gene_SNV_FNcountsboth.bed

paste GRCh38_mrg_full_gene_FNcounts.bed \
    GRCh38_mrg_full_gene_FNcountsIllonly.bed \
    GRCh38_mrg_full_gene_FNcountsboth.bed \
    GRCh38_mrg_full_gene_SNV_FNcounts.bed \
    GRCh38_mrg_full_gene_SNV_FNcountsIllonly.bed \
    GRCh38_mrg_full_gene_SNV_FNcountsboth.bed \
    GRCh38_mrg_full_gene_merged_FNcounts.txt
```

Where `GRCh38_mrg_full_gene.bed` is a list of all genes in GRCh38 (found in https://github.com/usnistgov/giab-cmrg-benchmarkset/blob/master/data/manually_created_files/GRCh38_mrg_full_gene.bed) and `patho_hifi_ilm.tsv` (**Supplemental File 2**) is a list of pathogenic variants with >90% likelihood of being missed by Hifi or Illumina as calculated by Stratomod.

#### Supplemental Note 2

*Predicting False Positives in PCR-free vs PCR-plus*

We asked if StratoMod could be used to predict where Illumina PCR-plus and PCR-free sequencing technologies have higher sequencing or mapping error rates that could produce false positive variant calls (**Supp Fig 5a**). This choice in comparison was motivated by the fact that PCR amplification is known to produce insertions and deletions in homopolymer repeats (stutter[1]) (**Supp Fig 5b**) and thus we hypothesized that the model would be able to precisely show the effect of homopolymer length on error rate, in addition to other repetitive genomic contexts.

We trained two models (for SNVs and INDELs) using Illumina PCR-free/plus VCF files. These models differed from those previously trained in “use-case 1” above in several ways. First, we used FP instead of FN as the error class, since in this case we were concerned with sequencing error modalities that could falsely give rise to variants. Second, we used all candidate sites (before filtering) from the DeepVariant VCFs since DeepVariant would likely filter out the sequencing errors we wished to interrogate. Third, we included DP and VAF as features in our model, bringing the total feature count up to 24 (see full list of features in **Supp Table 1**). For these models, we observed that both precision and recall (measured by AUC) were similar between HG004 and HG007 (with HG007 lagging slightly behind as expected given it was the holdout dataset) (**Supp Fig 5c, Supp Fig 6**). The negative class in the training sets for SNV and INDEL were 63% and 85% percent respectively (**Supp Table 2**).

Overall, the largest driving features (unsurprisingly) were VAF and DP as read from the input VCF files (**Supp Fig 6**), as many errors had low VAF and abnormally low or high DP (**Supp Fig 7**). However, other features with large effect included homopolymer length and homopolymer imperfect fraction (**Supp Fig 6a**). When observing the homopolymer length feature profiles directly from the model, we found that the likelihood of a FP generally increased with increasing length as expected from PCR stutter and sequencing biases. For INDELs, PCR-plus generally predicted more errors, with relatively small interactions between PCR and homopolymer length (**Supp Fig 5d**). Most homopolymers fell between the lengths of 0 to 50 or 0 to 15 bp for A/T and G/C homopolymers respectively, with the number of TPs and FPs decreasing exponentially with increasing length and ~100x more TPs in A/T than in G/C homopolymers longer than 10 bp (**Supp Fig 8**). Previous work had found that the number of FP INDELs in A/T homopolymers was much larger than in G/C homopolymers,[2] which was reflected in the feature rankings by Stratomod, but Stratomod also showed that G/C homopolymer length similarly predicts higher FP rates. Additionally, these feature plots provide more precise information regarding the length at which a certain relative error threshold will be crossed. In the case of SNVs, A homopolymers became more error-prone compared to the non-homopolymer baseline (the dotted lines in **Supp Fig 5d**) after 10 bp; G homopolymers of any length were more error prone (note that both A and G homopolymer profiles were similar to their complements, see **Supp Fig 8**). Increased SNV error rates in G/C homopolymers have been attributed to inhibition of base elongation in GC-rich regions during sequencing by synthesis[3] or to formation of non-B-DNA stem-loop motifs at G quadruplexes.[4] For INDELs, these thresholds were conditional on the sequencing technology, where PCR-free and PCR-plus were more error-prone than baseline after 13 and 11 bp respectively for A homopolymers. For G homopolymers this drop off occurred around 10 bp for both PCR-free and PCR-plus (note that in the case of C homopolymers these thresholds were 12 and 10 bp for PCR-free and PCR-plus respectively, see **Supp Fig 8**).

For INDELs, we also observed an unexpected increase from baseline (i.e., a higher likelihood of

TP vs. FP relative to non-homopolymers) in both A and G homopolymers for short lengths of between 4 and ~10 bp, with a peak around 8 bp. Because the EBM score is a function of both TP and FP rates, we hypothesized that the rate of true variants (TPs) increases faster than the rate of sequencing errors (FPs) in short homopolymer regions. Supporting this hypothesis, the ratio of TPs to FPs was higher for short homopolymers than for non-homopolymers, which may be caused by the higher rate of true INDEL variants in homopolymers (e.g., 39% of benchmark INDELs are in homopolymers 7 to 10 bp, while only 1.7% of the benchmark regions are in these homopolymer regions, **Supp Table 3**). Indeed, when plotting the TP and FP rates per base pairs covered by each homopolymer size, we saw that the TP rate increased faster than the FP rate for small homopolymers. As the homopolymer length increased, the FP/bp rate increased more than the TP/bp rate, which was reflected in the decreased EBM score. Interestingly, the TP/bp and FP/bp rates decreased for very large homopolymers, likely because large homopolymers are more likely to be excluded from the v4.2.1 GIAB benchmark, reflecting a limitation of the current training dataset (**Supp Fig 9**). These results explain the increase in EBM score for short homopolymers, followed by a decrease, before flattening out due to the small number of very long homopolymers included in the v4.2.1 benchmark. This deep-dive into a counter-intuitive result highlights both the challenges in interpreting the model's results, particularly that it is modeling the ratio of TPs to FPs rather than the FP rate per genomic bp, as well as its power in identifying unexpected associations of features with error rates.

We also noticed that the EBM INDEL scores had some sharp downward peaks for particular values of segmental duplication length and identity (**Supp Fig 10a**). When examining variants in the 2 largest peaks near 20 kbp, we found that they were caused by segmental duplication between chr7:142,450,000-142,526,000 on GRCh38 inside the T cell receptor beta locus. This region has a known issue in GRCh38, and the patch contains a ~20 kbp insertion, which is an extra tandem copy of the segmental duplication and causes many FP variant calls in Illumina and HiFi (**Supp Fig 10b**). It also intersects with 2 types of problematic reference regions identified in the recent T2T variants work: GRCh38 collapsed duplications and gnomAD inbreeding coefficient FPs, both of which annotate regions with FPs due to reads from extra copies of the region that are in most genomes but missing from the reference.[5] This result highlights a strength of this model to identify unexpected relationships between features and errors, and also suggests the possibility of adding new features associated with reference errors to future versions of the model.

These results indicate that these models can be used to precisely quantify FP error rates with respect to a meaningful, interpretable genomic context, which in turn could be useful in defining more accurate stratifications.

*Comparison of EBM performance to DeepVariant performance within candidate regions*

We next compared the accuracy of StratoMod's FP vs TP classifications to DeepVariant's, for the candidates generated by DeepVariant. We first examined the calibration of DeepVariant's genotype quality score (GQ) (**Supp Fig 11a**) and StratoMod's probability score (**Supp Fig 11b**). As expected from DeepVariant's richer information used for classification, it provided useful phred-based quality scores up to empirical scores of >50 (1 in 100,000 error rate) vs. the v4.2.1 benchmark, though it was somewhat overconfident for INDELs. StratoMod provides well-calibrated scores up to about 35 (1 in 3000 error rate) for SNVs and 25 (1 in 300 error rate) for INDELs. In **Supp Figure 11b**, we assign a label on "1" to any StratoMod probability > .5 and "0" otherwise. We then made a Venn diagram showing intersections between DeepVariant and StratoMod predicted TP labels against the benchmark when combining SNVs and indels for PCR-free and PCR-plus Illumina. 99.4% of the benchmark variants were classified as TPs by both StratoMod and DeepVariant. Of the remaining 0.6%, most (36,675) were classified incorrectly as FPs by StratoMod and as TPs by DeepVariant, whereas 1,765 were correctly classified as TPs by StratoMod and not DeepVariant. In addition, StratoMod incorrectly classified 44,629 variants as TPs that were correctly filtered by DeepVariant, many more than the 3,930 uniquely incorrect TPs in DeepVariant. When intersecting these uniquely mis-classified variants with features, DeepVariant uniformly performed better, but ~25% of false variants uniquely classified as TPs by DeepVariant were in regions difficult to map with 250 bp reads, suggesting DeepVariant may benefit from additional genome mappability features.

*Comparison of EBMs to other commonly used models*

We compared the performance of an EBM model to XGBoost (XGB), Random Forest (RF), Logistic Regression (LR), and Decision Tree (DT) models in classifying variants as errors using the same training and testing data. In all cases, performance was comparable when examining the area under the curve (AUC) for the receiver operator characteristic (ROC) curve and precision-recall (PR) curves (**Supp Table 4**). Note that the RF SNV model did not complete within the constraints imposed by our compute cluster.

While similar, StratoMod performed slightly better than both DT and LR models and slightly worse than RF and XGB models for both SNVs and INDELs. This is not surprising considering that RF and XGB are much more flexible than EBMs, and EBMs in turn are more flexible than LR models. DT models might be more flexible than EBMs given that they have less restrictions when building trees, but DT models also tend to be brittle as they are not ensemble models (unlike EBMs). Notably, the models that did perform slightly better than EBMs were also blackbox models (e.g. unable to be inspected analogously to EBMs).

These data demonstrated that for this use case, using EBMs for the classification algorithm in StratoMod performed similarly to other commonly used models. Slight performance benefits may be had with blackbox models such as random forest or XGBoost, albeit with a loss of interpretability and potentially much higher compute requirements.

#### Supplemental figures:

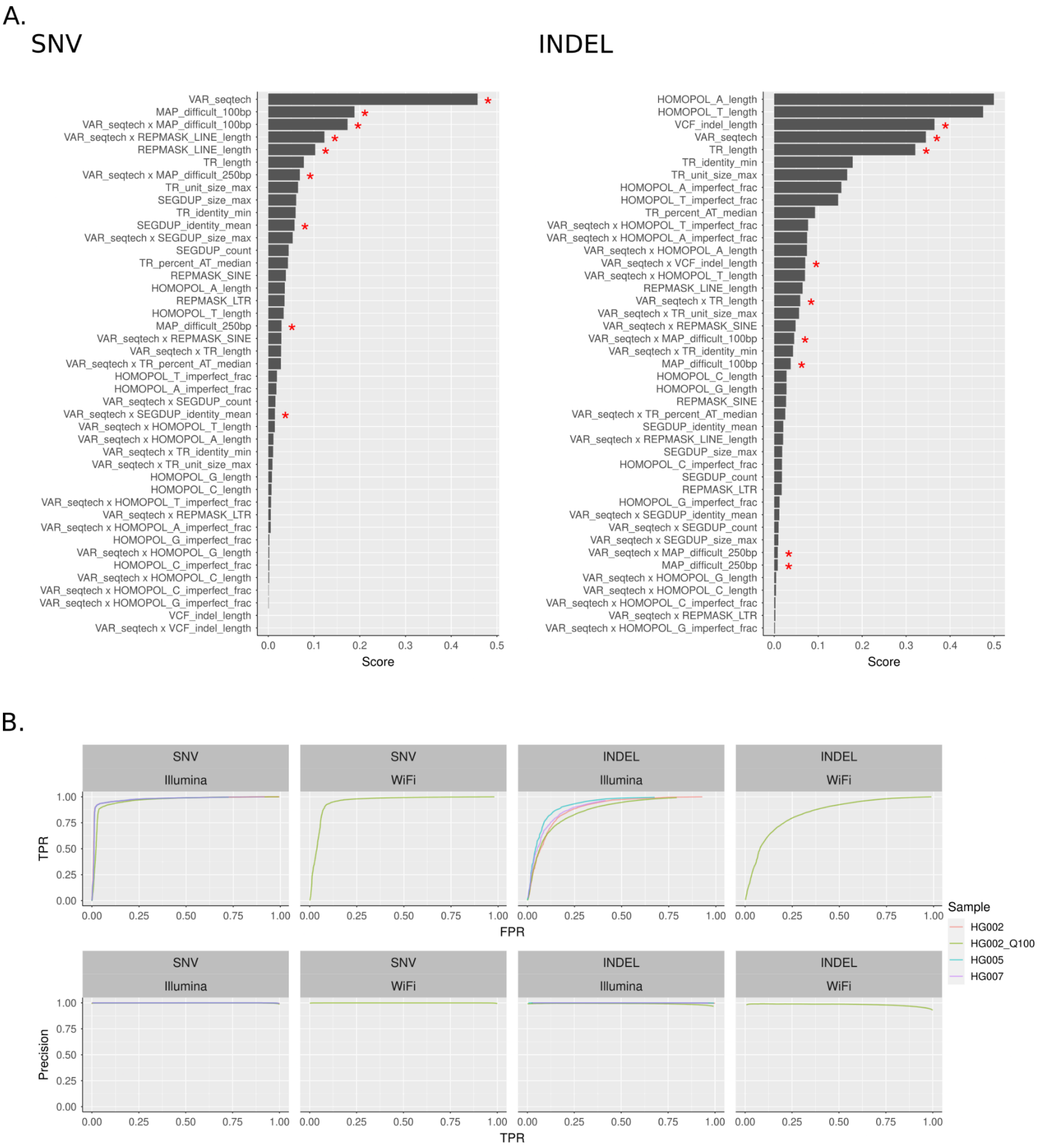


**Supplemental Figure 1**: Global importance plots (a) and performance curves (b) for Hifi vs Illumina Clinvar experiment. TPR = true positive rate. Train was done on 80% of “HG002_Q100” and test was done on 20% of “HG002_Q100” and the whole of the other genomes, which were all in terms of the v4.2.1 Genome in a bottle benchmark.

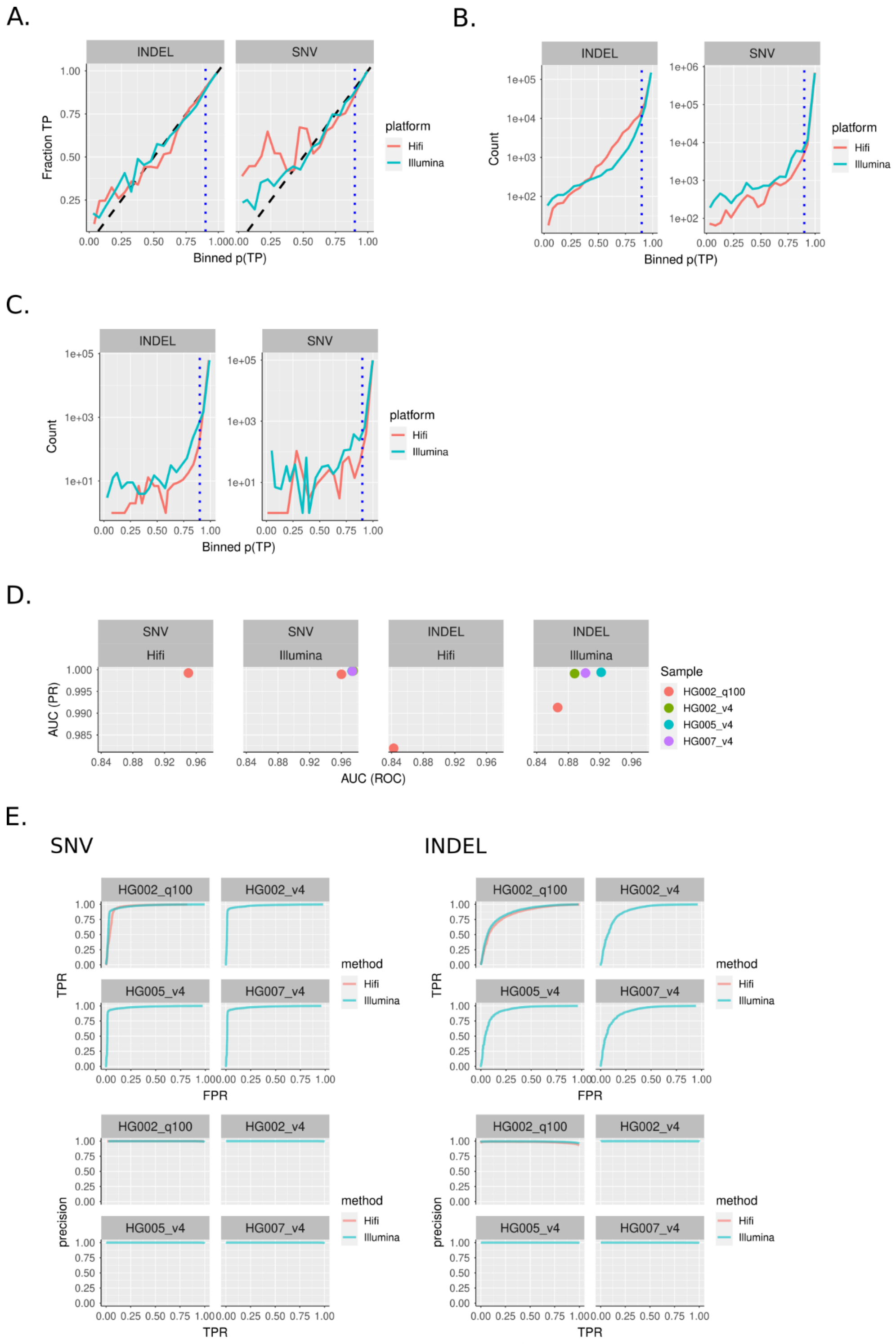


**Supplemental Figure 2**: Performance metrics for FN Hifi vs Illumina model. A) calibration curves where the dotted black line is perfect calibration and blue dotted line is the 90% threshold used in the analysis. B-C) Counts for each bin showed in (A) for test data set (B) and clinvar variants (C) D) AUC under precision-recall (PR) and receiver-operator (ROC) curves for all models. D) Raw ROC and PR curves for each genome and platform.

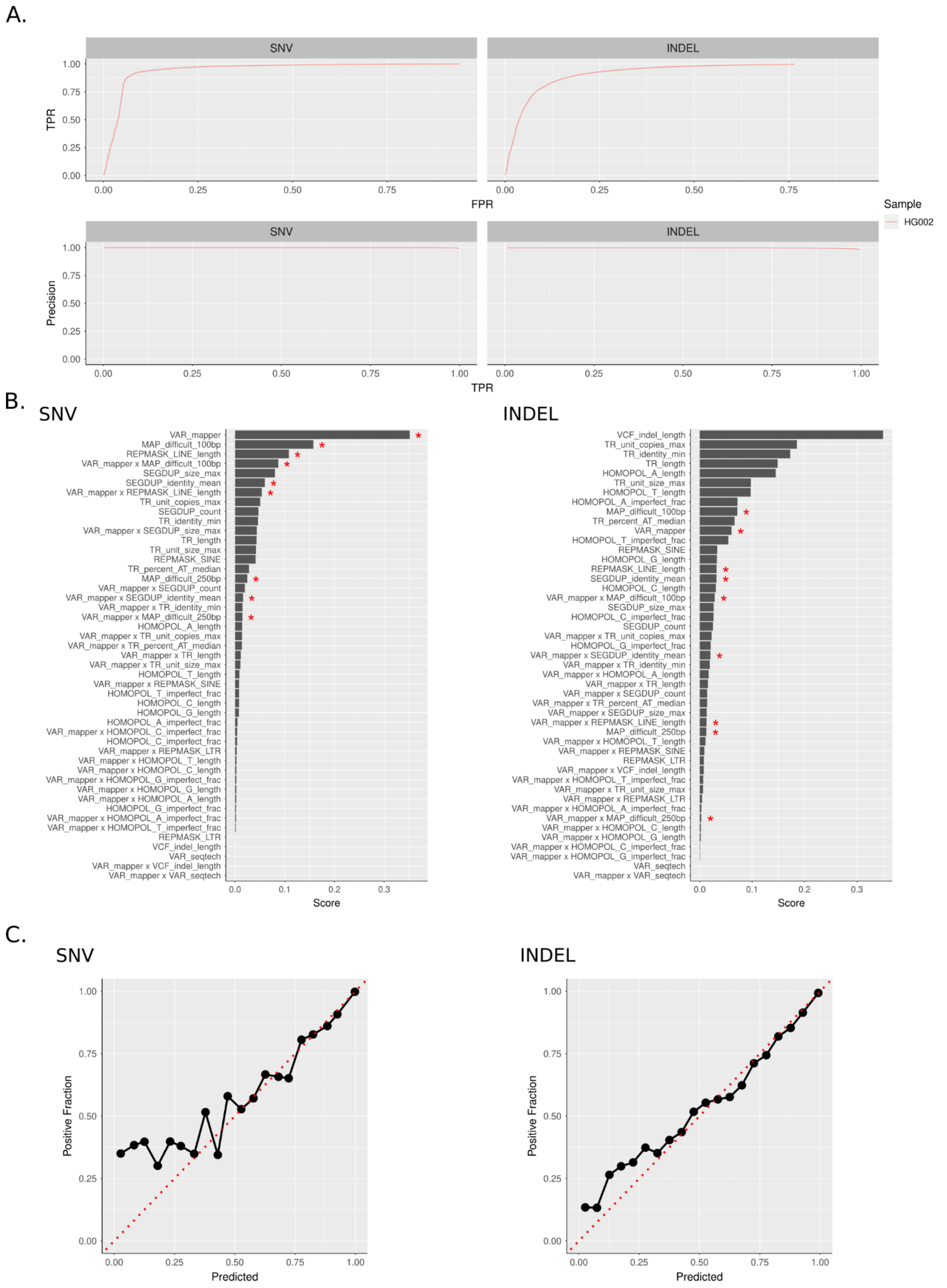


**Supplemental Figure 3:** Performance metrics for Element VG vs BWA FN model a) ROC and PR curves b) feature importance plots c) calibration curves (dotted line is perfect calibration).

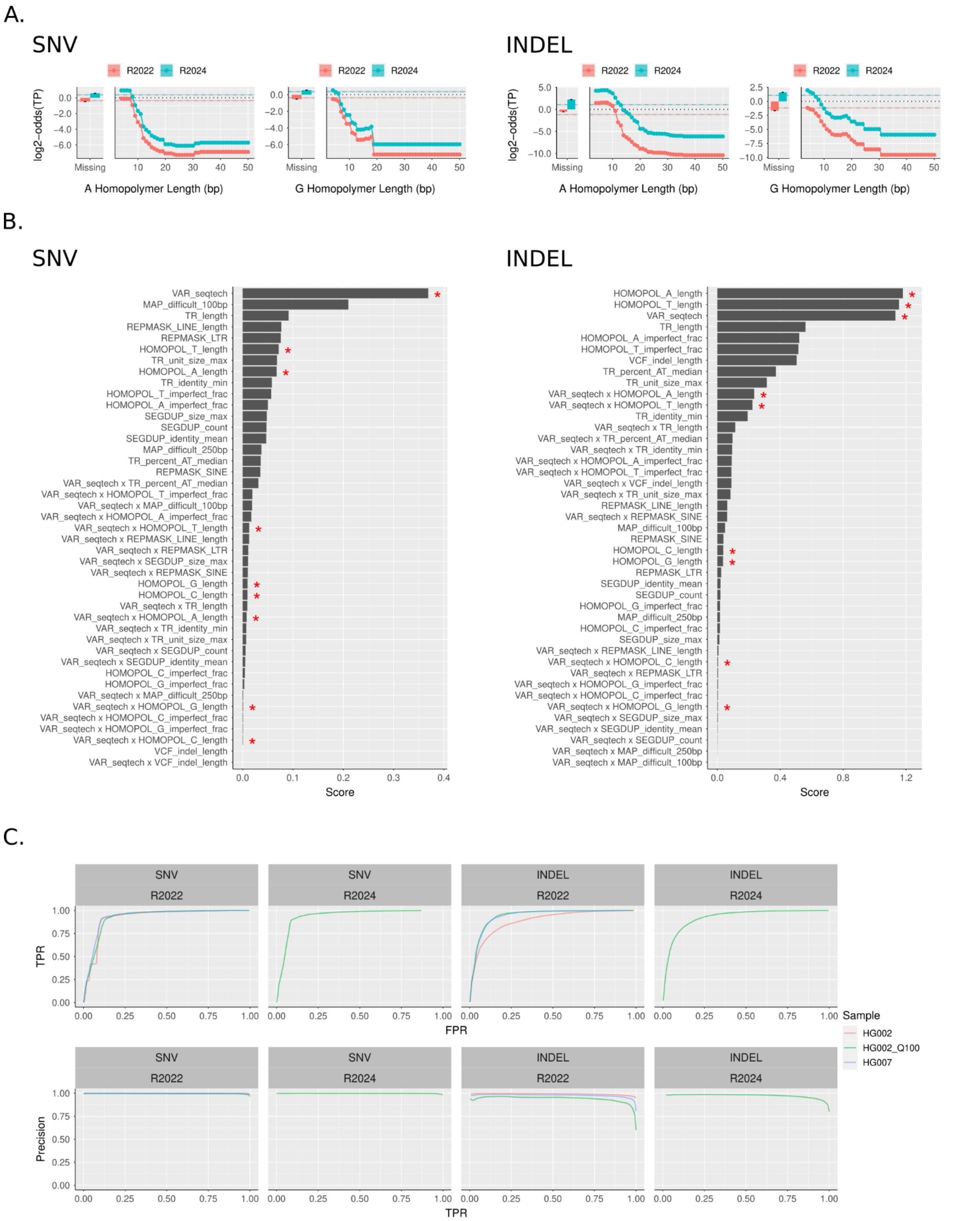


**Supplemental Figure 4**: StratoMod utilized to compare different versions of emerging technologies (Ultima versions 2022 vs 2024) in terms of their ability to detect FN errors. a) homopolymer profiles for INDELs and SNVs b/t old and new b) global feature plots for models and c) performance curves for each model

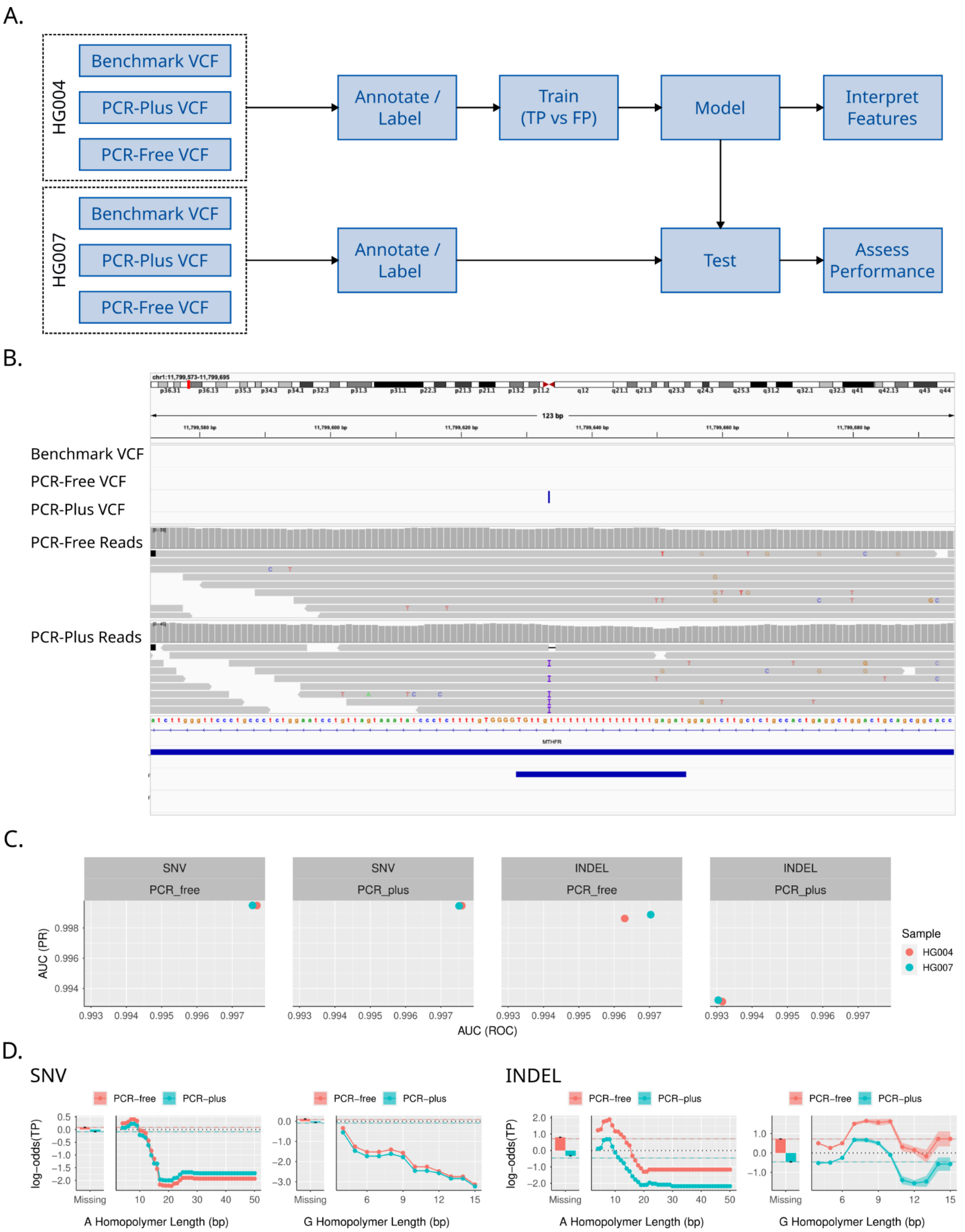


**Supplemental Figure 5**: StratoMod revealed context-specific regions where false positives are likely to occur and show relative performance of PCR-free and PCR-plus technologies. a) Overview of experimental setup. Two VCFs from PCR-free and PCR-plus were compared to the GIAB benchmark, concatenated, and annotated before fitting SNVs and INDELs in the EBM framework. HG005 was annotated and used to test the EBM model. b) IGV session depicting a false positive call identified by this model c) performance characteristics of HG004 (train) and HG007 (test). d) EBM plots showing A and G homopolymer profiles. The x axis in all plots was truncated to only show homopolymers <50bp and <15bp for A and G respectively. The bar plots on the left of each plot show the value for non-homopolymers (“missing”). The y axis in the top row is the log odds of a TP outcome. Each colored dotted line in the top row is the “baseline” error rate for the corresponding sequencing technology. The bottom plots show the distribution of homopolymer lengths (TPs are above the dotted line and FPs are below).

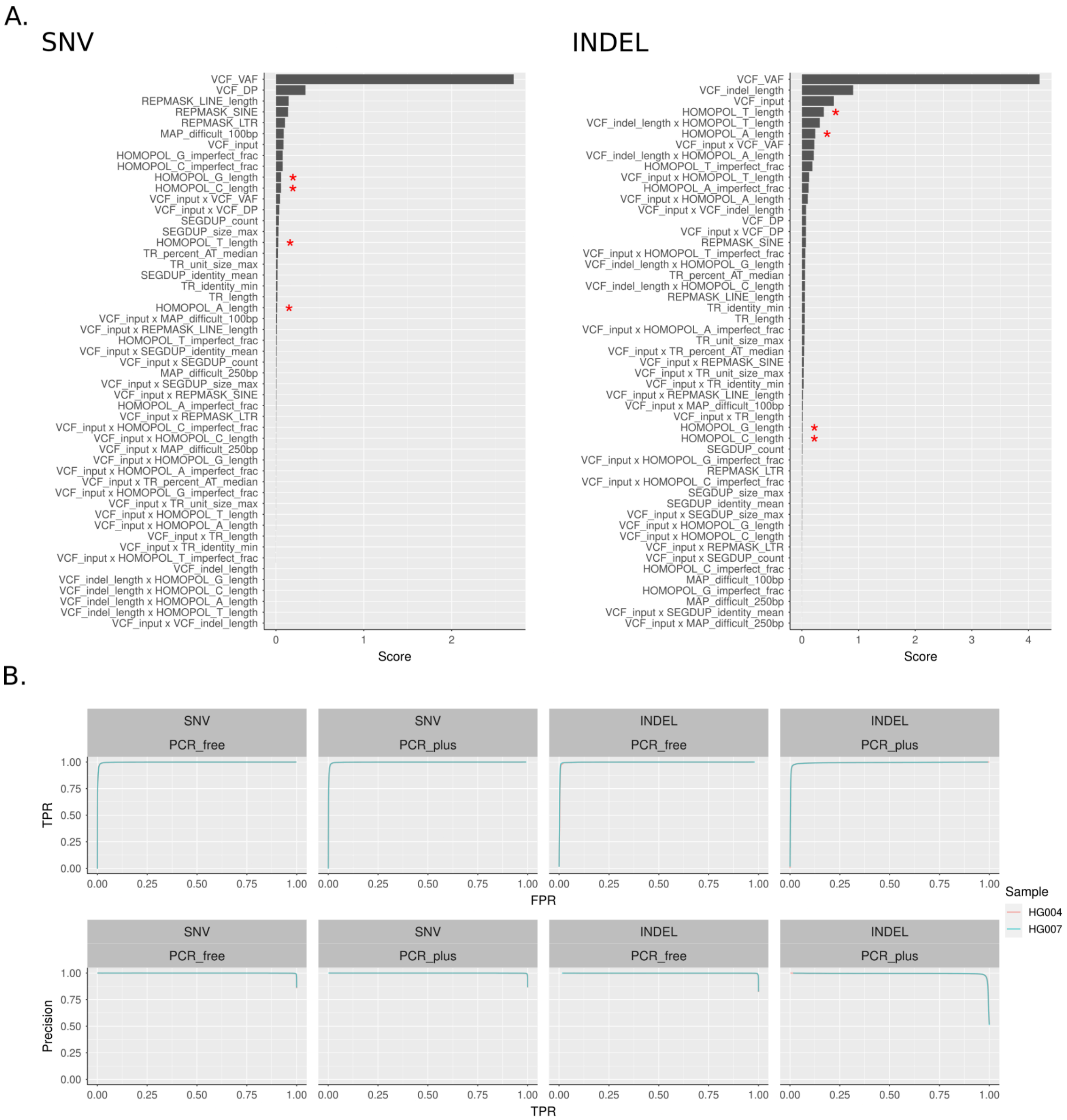


**Supplemental Figure 6**: Performance of PCR-free/plus models. a) global feature plots and b) ROC and PR curves

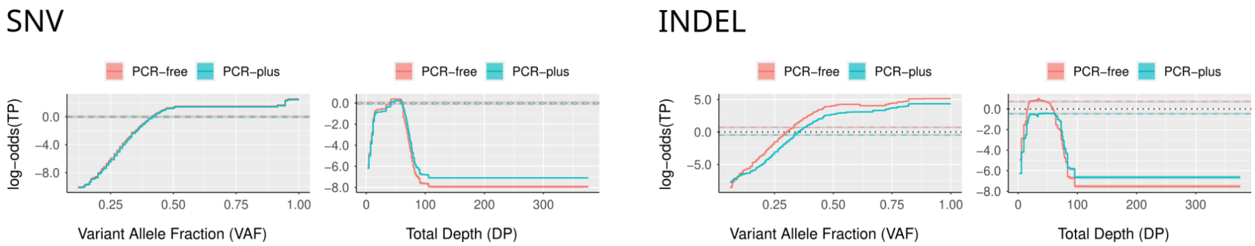


**Supplemental Figure 7**: VAF and DP feature plots for both SNV and INDEL FP models.

#### SNV

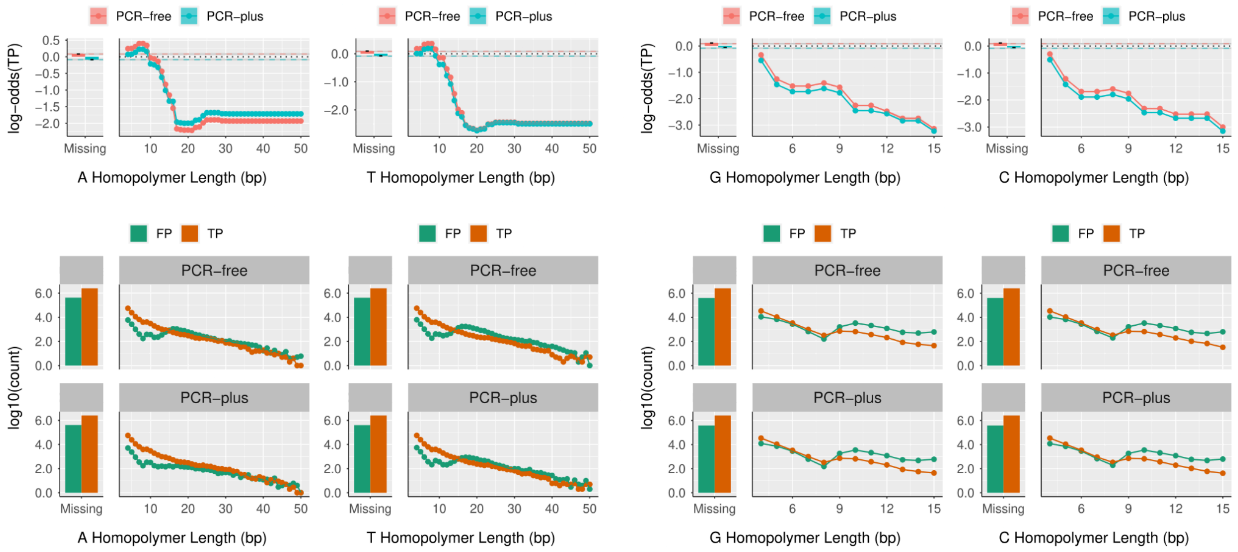


#### INDEL

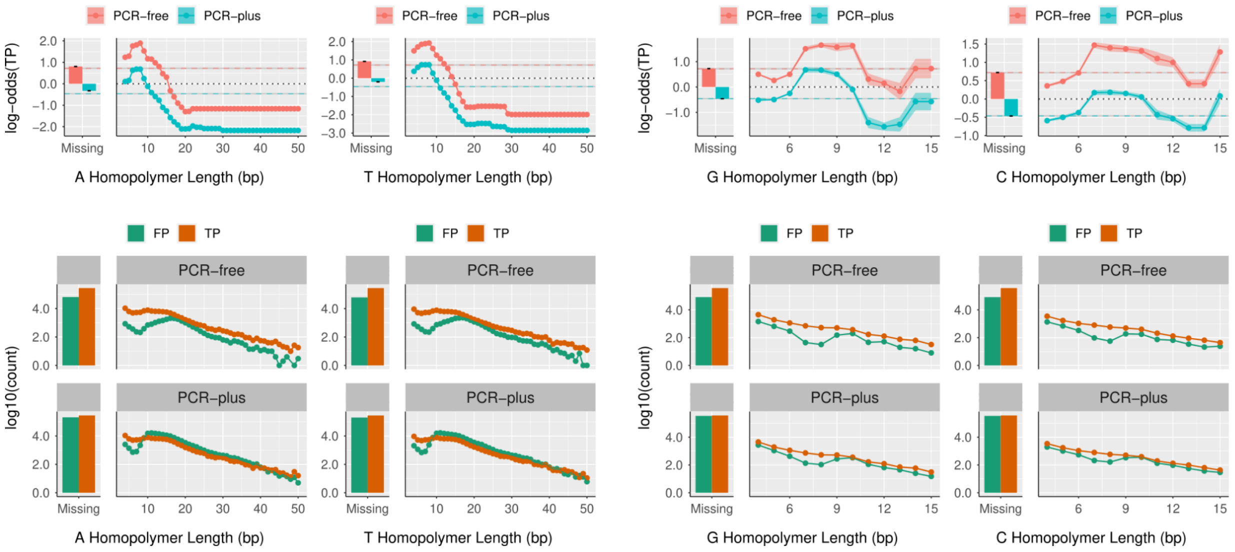


**Supplemental Figure 8**: Complementary homopolymers for the FP model for both SNVs and INDELs.

A.

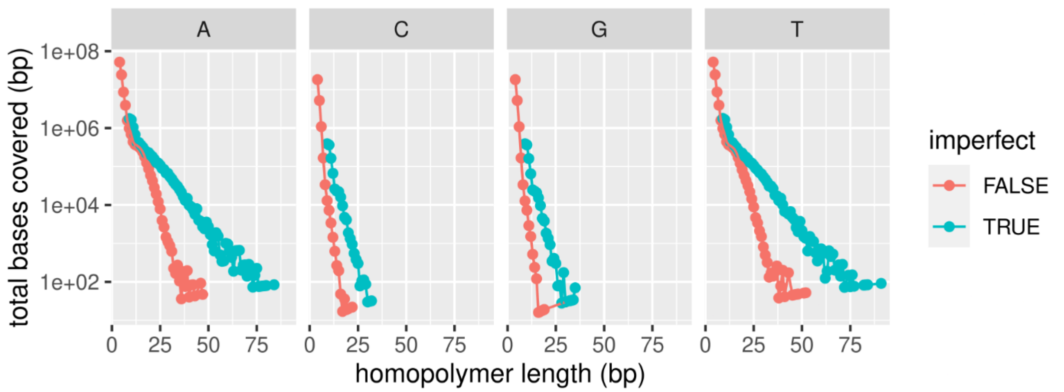


B.

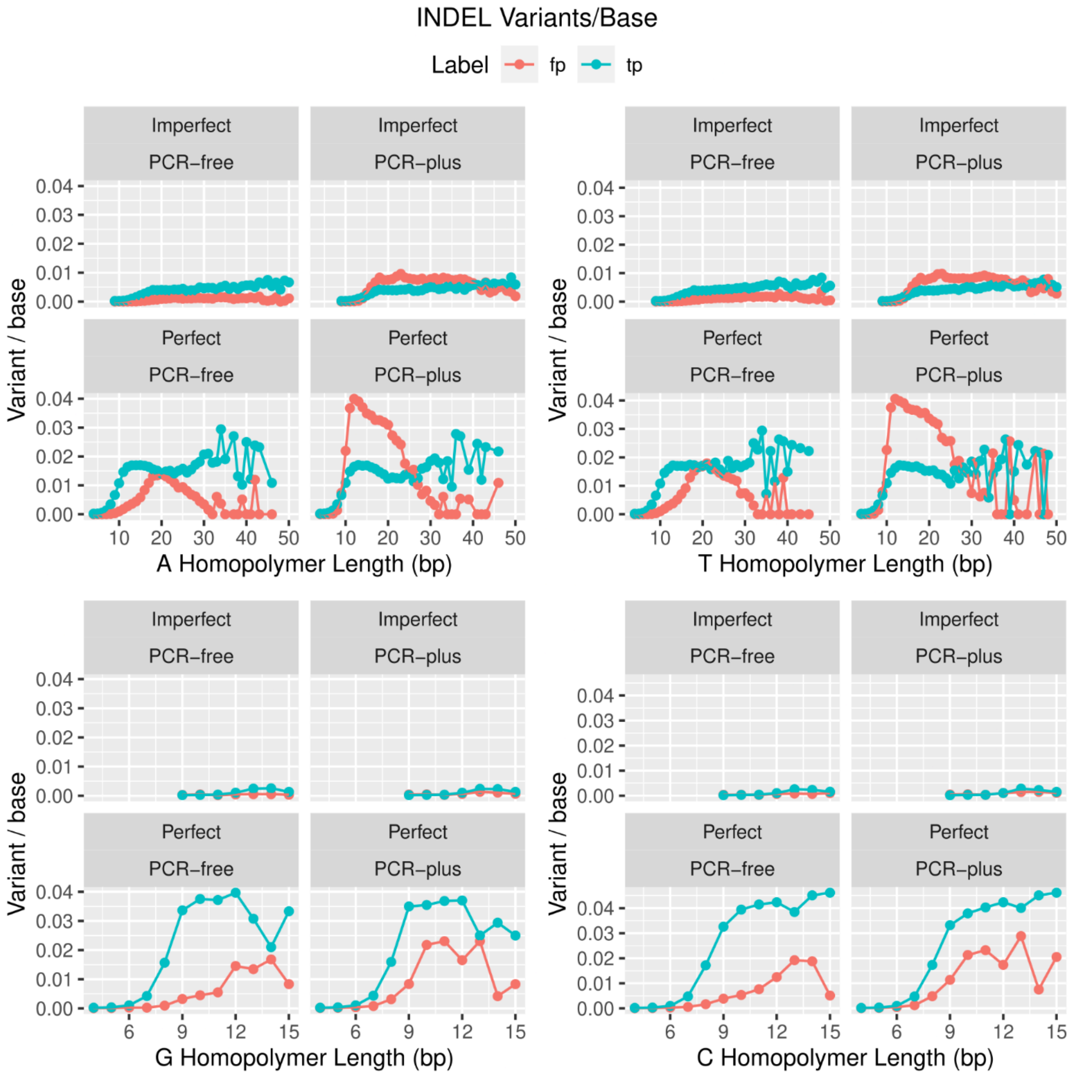


**Supplemental Figure 9**: Decomposition of homopolymer features for INDELs to better understand the EBM scores. A) total bases covered for each base homopolymer stratified by length B) number of variants normalized to total bases covered (from A) for each base stratified by length

A.

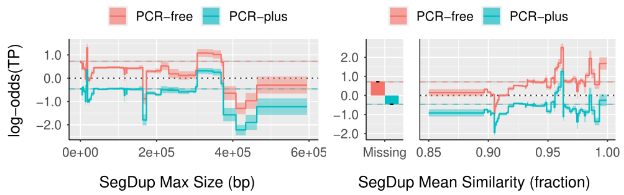


B.

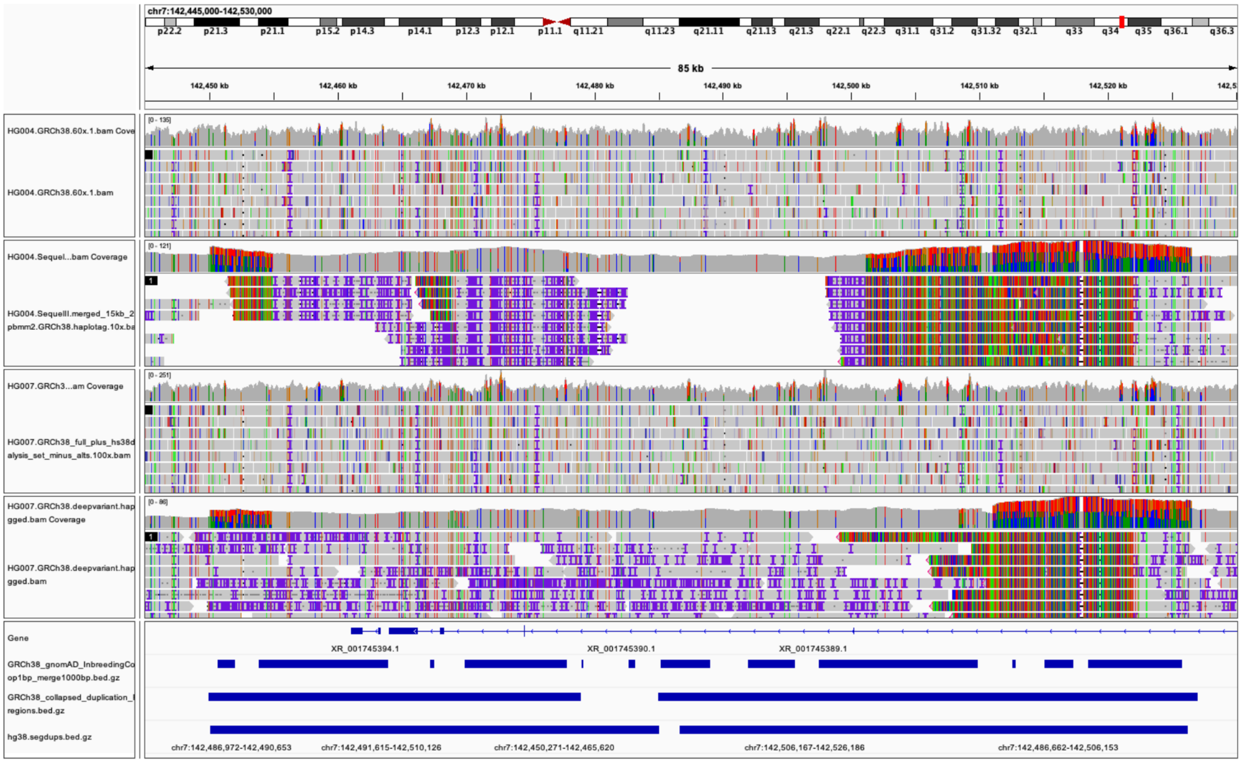


**Supplemental Figure 10:** Explainable segdup features facilitated identification of mapping errors in known hard regions. A) The INDEL feature profiles for the FP model for segdup identity and length (note the peak at ~20k). B) IGV screenshot showing FPs due to mapping errors from a duplicated region in HG004/HG007 due to an error in GRCh38 missing a copy of genes in this T cell receptor beta locus.

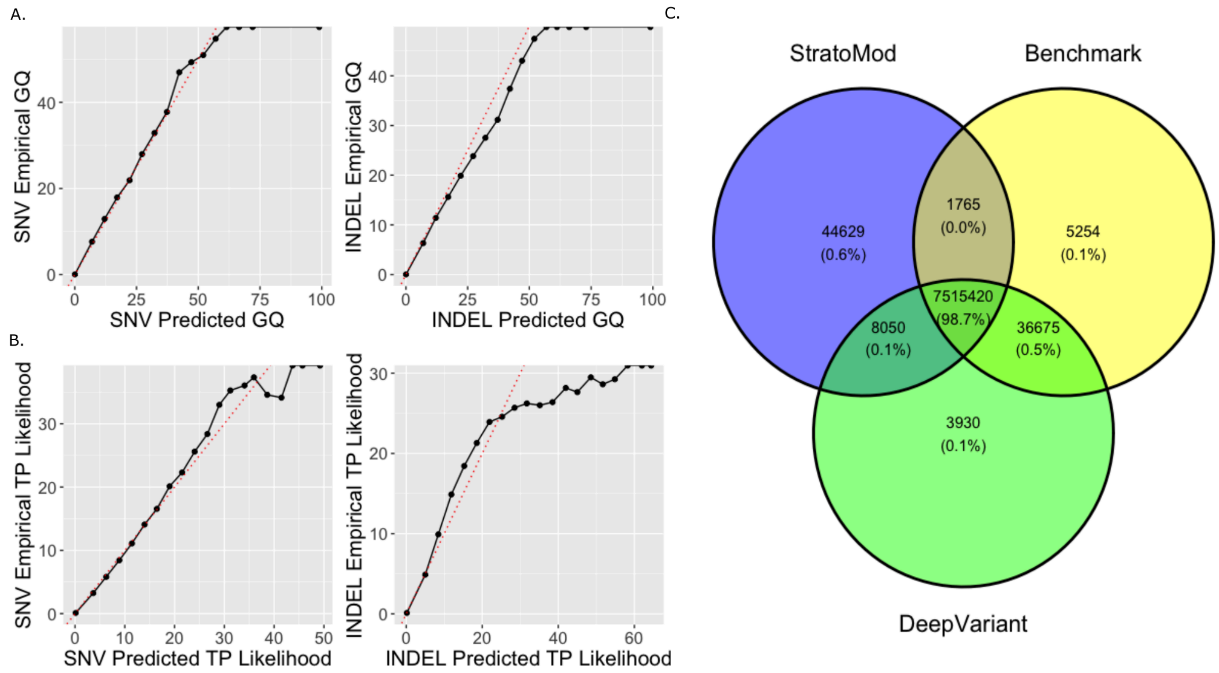


**Supplemental Figure 11**: Comparison of Stratomod calibration and accuracy relative to deep learning-based method DeepVariant. a) Predicted genotype quality score (GQ) from DeepVariant plotted against an empirically derived GQ measure, -10*log10(FP/(FP+TP)), using the GIAB v4.2.1 small variant benchmark. b) PHRED-scaled plots depicting calibration for how well StratoMod confidently predicts TPs (or values near 1 for its predictions) similar to a typical genotype quality score c) Comparison of StratoMod and DeepVariant performance against the benchmark values for the candidate sites from DeepVariant used in our model.

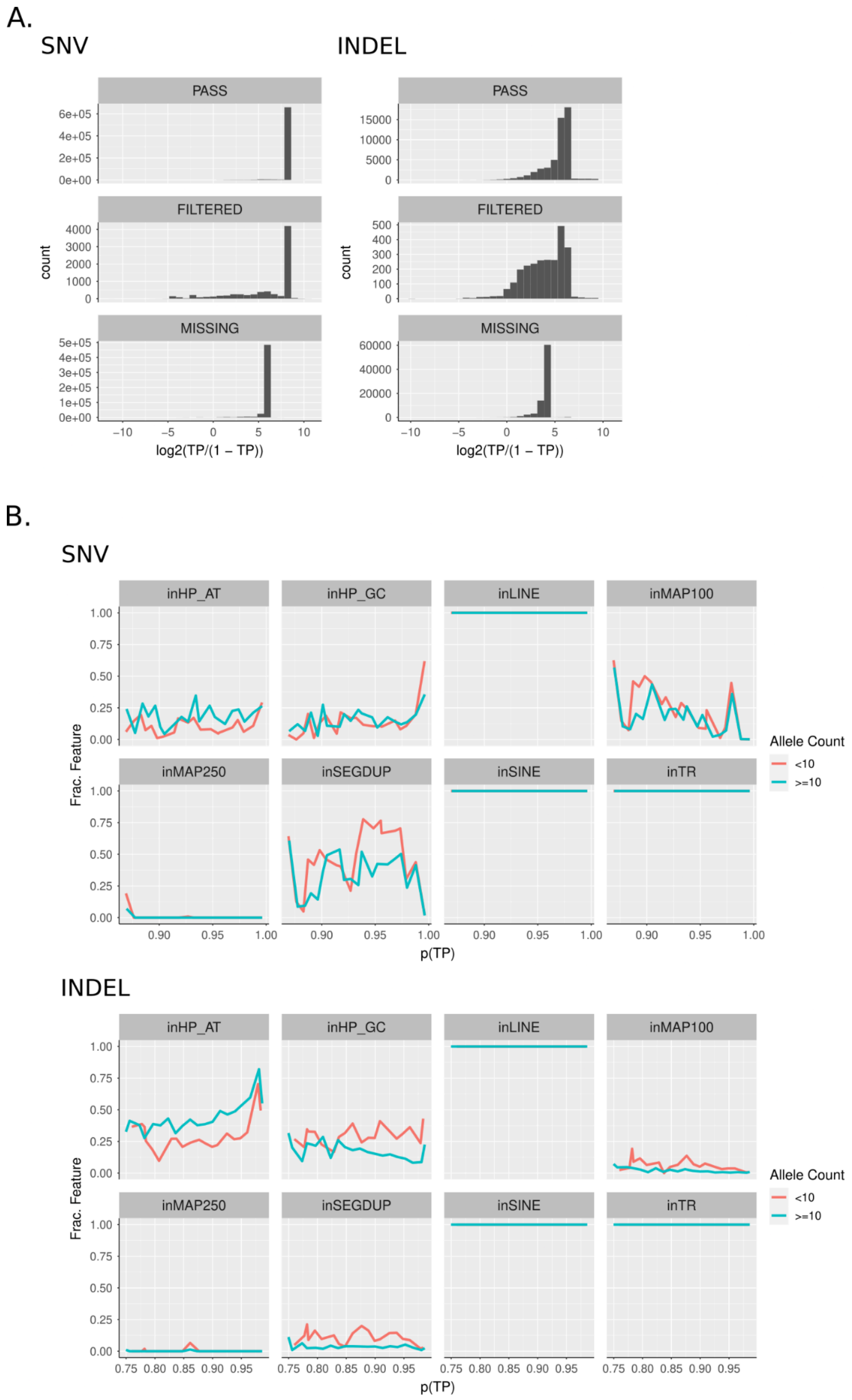


**Supplemental Figure 12**: a) Histograms of Clinvar variants that either corresponded to a PASS gnomAD variant ("PASS"), filtered gnomAD variant ("FILTERED") or no gnomAD variant ("MISSING"). b) Variants binned by probability and stratified by region type and allele count. Y axis is fraction of variants per bin that are within the denoted region type.

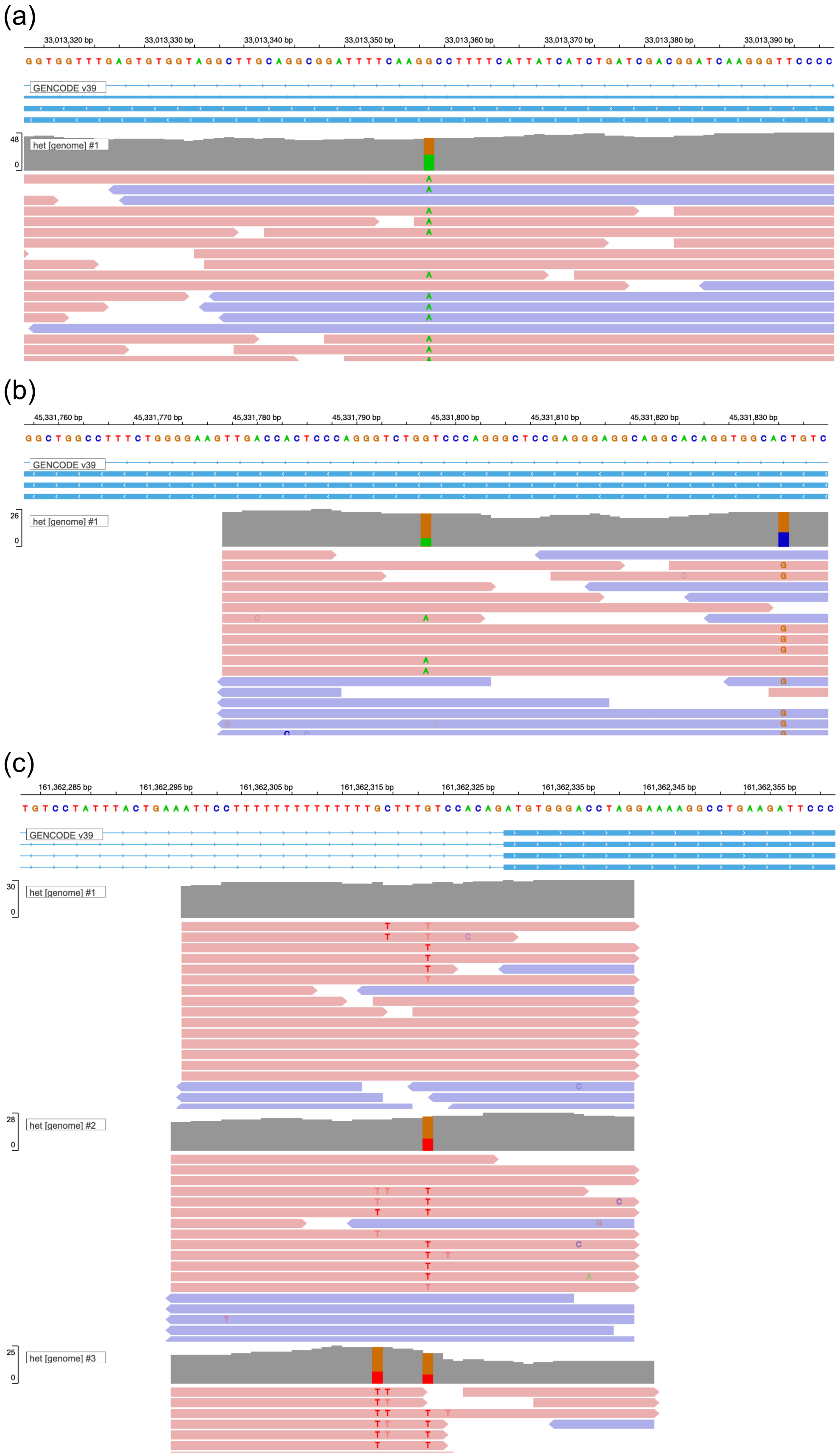

(a)
33,013,320 bp
33,013,330 bp
33,013,340 bp
33,013,350 bp
33,013,360 bp
33,013,370 bp
33,013,380 bp
33,013,390 bp
GENCODE v39
48
0
het [genome] #1
(b)
45,331,760 bp
45,331,770 bp
45,331,780 bp
45,331,790 bp
45,331,800 bp
45,331,810 bp
45,331,820 bp
45,331,830 bp
GENCODE v39
26
0
het [genome] #1
(c)
161,362,285 bp
161,362,295 bp
161,362,305 bp
161,362,315 bp
161,362,325 bp
161,362,335 bp
161,362,345 bp
161,362,355 bp
GENCODE v39
30
0
het [genome] #1
28
0
het [genome] #2
25
0
het [genome] #3

(d)

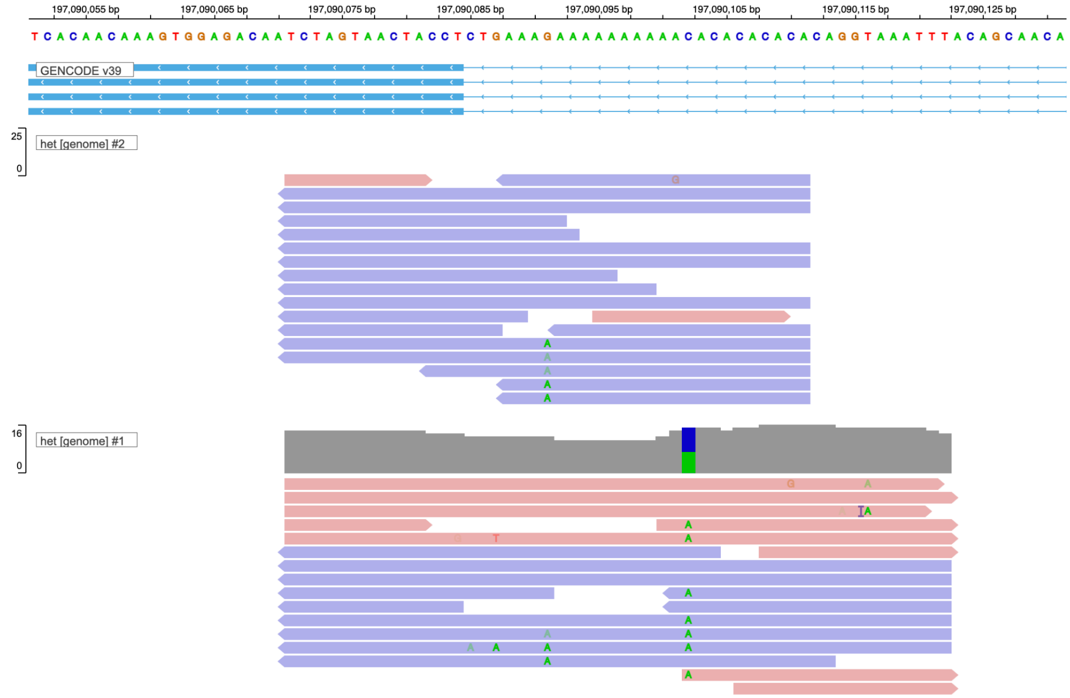


**Supplemental Figure 13**: Examples of gnomAD variants on GRCh38 chr1 that were filtered despite having a low error rate predicted by Stratomod (score>0.99). Upon curation, these generally fell in two categories: (a-b) variants in a very small number of samples, typically one or two, that appear likely to be true and were likely incorrectly filtered by gnomAD. (c-d) variants with evidence of systematic errors indicated by strand bias in homopolymers, and were generally called in more samples than the first category but still generally <100.

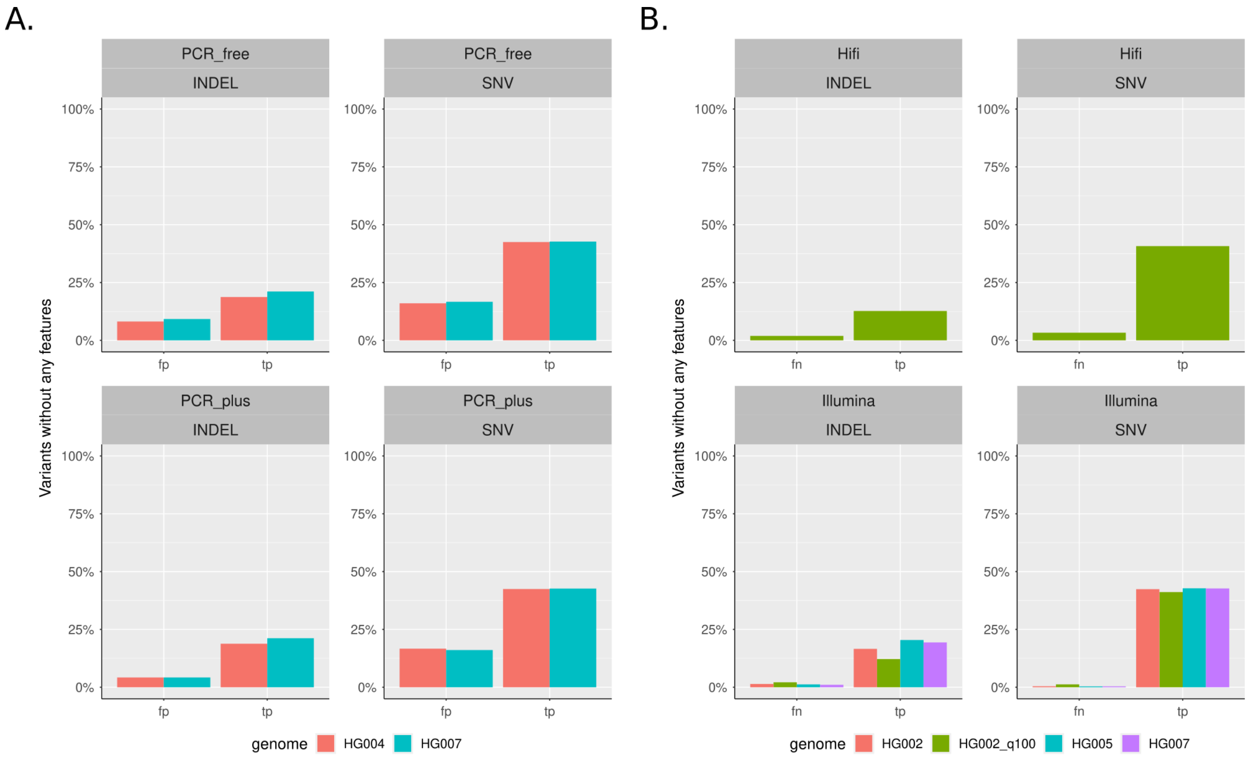


**Supplemental Figure 14**: Percent of variants which have no coverage by our engineered feature set in the a) Illumina PCR-free/plus FP model and b) the HiFi/Illumina FN model. Note that DP and VAF were excluded from this analysis, as they are expected to cover all FPs and TPs and no FNs.

### Supplemental Tables

**Supplemental Table 1**: Overview of features used in each model

| Feature Name | Description | Domain | Models |
|---|---|---|---|
| VCF_input | The VCF file corresponding to the origin of the variant; used to represent the different technologies/pipelines (eg PCR-free vs PCR-plus) | categorica l | all |
| VCF_VAF | The VAF (Variant Allele Fraction) value as recorded in the VCF file | [0-1] | PCR-free /plus only |
| VCF_DP | The DP (Depth of Coverage) value as recorded in the VCF file | [1-inf]; integer | PCR-free /plus only |
| VCF_indel_length | The length of an INDEL (0 in the case of SNVs) | [-inf-inf]; integer | all |
| HOMOPOL_<base>_length | The length of a homopolymer region (where <base> is either A, T, G, or C) | [1-inf]; integer | all |
| HOMOPOL_<base>_imperfect_frac | The fraction of bases in a homopolymer that are not <base> | [0-1] | all |
| TR_length | The length of a tandem repeat region | [1-inf]; integer | all |
| TR_unit_size_max | The size of a single repeat unit that is repeated TR_unit_copies times in a tandem repeat region. For overlapping regions the maximum was used. | [1-inf]; integer | all |
| TR_identity_min | The similarity of each unit across the tandem repeat (corresponds to ‘perMatch’ in the TRF-based UCSC simple repeats database). For overlapping regions the minimum was used. | [1-inf]; | all |
| TR_percent_AT_median | The percentage of the tandem repeat region which is an A or T base | [0-100]; integer | all |
| REPMASK_SINE | TRUE is the variant intersects with a SINE | boolean | all |
| REPMASK_LTR | TRUE if the variant intersects with an LTR | boolean | all |
| REPMASK_LINE_length | The length of the LINE in which this | [1-inf]; | all |

| | | | |
|---|---|---|---|
| | variant is found | integer | |
| MAP_difficult_Xbp | TRUE if the variant intersects with a hard-to-map region for read pairs of length X (where X is either 100 or 250). | boolean | all |
| SEGDUP_size_max | The length of the segmental duplication region. For overlapping regions, the maximum was used. | [1000-inf]; integer | all |
| SEGDUP_identity_mean | The similarity of this segmental duplication to others (corresponds to “fracMatchIndel” from the genomic super dups database). For overlapping regions the mean was used. | [0-1] | all |
| SEGDUP_count | The number of segmental duplications overlapping this region. | [1-inf]; integer | all |

**Supplemental Table 2**: Model training summary

| N Rows | Percent TP | Variant Type | Subsets | Error Type |
|---|---|---|---|---|
| 2939508 | 94.4 | INDEL | Hifi = 1452626; Illumina = 1486882 | fn |
| 11412592 | 98.83 | SNV | Hifi = 5718674; Illumina = 5693918 | fn |
| 2406562 | 97.62 | INDEL | VG = 1202820; BWA = 1203742 | fn |
| 11407256 | 99.24 | SNV | VG = 5704070; BWA = 5703186 | fn |
| 2640324 | 70.16 | INDEL | R2024 = 1366776; R2022 = 1273548 | fn |
| 11388272 | 97.11 | SNV | R2024 = 5689340; R2022 = 5698932 | fn |
| 2375368* | 63.69 | INDEL | PCR-free = 929278; PCR-plus = 1446090 | fp |
| 12311190* | 85.95 | SNV | PCR-free = 6177338; PCR-plus = 6133852 | fp |
| 383191 | 92.45 | INDEL | NA | fp+fn |
| 1429350 | 98.06 | SNV | NA | fp+fn |

* All used the GIAB assembly based small variant benchmark from the T2T-HG002-Q100v0.9 assembly aligned to GRCh38 under https://ftp-trace.ncbi.nlm.nih.gov/ReferenceSamples/giab/data/AshkenazimTrio/analysis/NIST_HG002_DraftBenchmark_defrabbV0.011-20230725/ except for these, which used the GIAB v4.2.1 small variant benchmark from https://ftp-trace.ncbi.nlm.nih.gov/ReferenceSamples/giab/release/AshkenazimTrio/HG002_NA24385_son/NISTv4.2.1/GRCh38/

**Supplemental Table 3**: Homopolymer TP likelihood increase for length 7-10bp

| **Homopolymer length** | **Number of 1 bp INDELs** | **Number of bp covered by homopolymer + 5 bp slop on each side** |
|---|---|---|
| 4 to 6 bp | 66134 | 747,661,356 |
| 7 to 11 bp | 80261 | 47,824,121 |
| >11 bp | 31624 | 17,659,926 |
| All benchmark regions | 204853 | 2,765,733,593 |

**Supplemental Table 4**: Comparison of EBM performance to those of other popular machine-learning methods. OOM: out of memory

| Variant | Metric | StratoMod | DT | LR | RF | XGB |
|---|---|---|---|---|---|---|
| INDEL | ROC | 99.5% | 99.2% | 99.2% | 99.8% | 99.8% |
| INDEL | PR | 99.7% | 99.4% | 99.5% | 99.9% | 99.9% |
| SNV | ROC | 99.8% | 99.4% | 99.5% | OOM | 99.9% |
| SNV | PR | 99.9% | 99.8% | 99.9% | OOM | 100.0% |

**Supplemental Table 5**: Software packages and versions

| **Tool/package name** | **Version** |
|---|---|
| rtg-tools (vcfeval) | 3.12.1 |
| bedtools | 2.30.0 |
| Interpretml (the EBM python package) | 0.2.7 |
| samtools | 1.14 |
| bcftools | 1.19 |

**Supplemental Table 6:** VCF files used throughout analysis (all with GRCh38 as reference)

| Description | Sample | Coverage (X) | Source |
|---|---|---|---|
| Illumina PCR-Free | HG002 | 40 | https://storage.googleapis.com/brain-genomics-public/research/sequencing/grch38/vcf/hiseqx/wgs_pcr_free/40x/HG002.hiseqx.pcr-free.40x.deepvariant-v1.0.grch38.vcf.gz |
| Illumina PCR-Free | HG004 | 40 | https://storage.googleapis.com/brain-genomics-public/research/sequencing/grch38/vcf/hiseqx/wgs_pcr_free/40x/HG004.hiseqx.pcr-free.40x.deepvariant-v1.0.grch38.vcf.gz |
| Illumina PCR-Plus | HG004 | 40 | https://storage.googleapis.com/brain-genomics-public/research/sequencing/grch38/vcf/hiseqx/wgs_pcr_plus/40x/HG004.hiseqx.pcr-plus.40x.deepvariant-v1.0.grch38.vcf.gz |
| Illumina PCR-Free | HG005 | 40 | https://storage.googleapis.com/brain-genomics-public/research/sequencing/grch38/vcf/hiseqx/wgs_pcr_free/40x/HG005.hiseqx.pcr-free.40x.deepvariant-v1.0.grch38.vcf.gz |
| Illumina PCR-Free | HG007 | 40 | https://storage.googleapis.com/brain-genomics-public/rese |

| | | | |
|---|---|---|---|
| | | | arch/sequencing/grch38/vcf/hiseqx/wgs_pcr_free/40x/HG007.hiseqx.pcr-free.40x.deepvariant-v1.0.grch38.vcf.gz |
| Illumina PCR-Plus | HG007 | 40 | https://storage.googleapis.com/brain-genomics-public/research/sequencing/grch38/vcf/hiseqx/wgs_pcr_plus/40x/HG007.hiseqx.pcr-plus.40x.deepvariant-v1.0.grch38.vcf.gz |
| PacBio HiFi | HG002 | 37 | https://ftp-trace.ncbi.nlm.nih.gov/giab/ftp/data/AshkenazimTrio/analysis/PacBio_CCS_15kb_20kb_chemistry2_10312019/GRCh38/deepvariant_HG002_GRCh38_15kb_37X_SequelII.vcf.gz |
| PacBio HiFi | HG005 | | https://ftp-trace.ncbi.nlm.nih.gov/ReferenceSamples/giab/release/ChineseTrio/HG005_NA24631_son/NISTv4.2.1/GRCh38/SupplementaryFiles/inputvcfsandbeds/HG005_GRCh38_1_22_PacBio_HiFi_DeepVariant.vcf.gz |
| PacBio HiFi | HG007 | 21 | https://storage.googleapis.com/brain-genomics-public/research/sequencing/grch38/vcf/pacbio_hifi |

| | | | |
|---|---|---|---|
| | | | /HG007.pacbio-hifi.21x.deepvariant-v1.0.grch38.vcf.gz |
| ClinVar | n/a | n/a | https://ftp.ncbi.nlm.nih.gov/pub/clinvar/vcf_GRCh38/archive_2.0/2022/clinvar_20220812.vcf.gz |
| Ultima R2024 | HG002 | 40 | https://giab-data.s3.amazonaws.com/ultima-GIAB-Feb-2024/DeepVariant_vcfs/NA24385-Z0027.annotated.AF.vcf.gz |
| Ultima R2022 | HG002 | 40 | https://s3.amazonaws.com/ultima-selected-1k-genomes-vcf-only/DeepVariant_vcfs/HG002_005401-UGAv3-1-CACATCCTGCATGTGAT.vcf.gz |
| Ultima R2022 | HG007 | 40 | https://s3.amazonaws.com/ultima-selected-1k-genomes-vcf-only/DeepVariant_vcfs/HG007_004731-UGAv3-33-CATGCAGCGCTAATGA.vcf.gz |
